# Reduced activity of nucleus accumbens parvalbumin-expressing fast-spiking inhibitory neurons causes convulsive seizures

**DOI:** 10.64898/2026.03.09.710428

**Authors:** Toshimitsu Suzuki, Takahiko Kondo, Tetsushi Yamagata, Yurina Hibi, Hiroaki Mizukami, Kenta Kobayashi, Kazuhiro Yamakawa

**Author notes:** Address correspondence to: Toshimitsu Suzuki, Ph.D. and Kazuhiro Yamakawa, Ph.D. Department of Neurodevelopmental Disorder Genetics, Institute of Brain Science, Nagoya City University Graduate School of Medical Sciences, Nagoya, 467–8601, Japan; or.

## Abstract

Pathogenic mutations in *STXBP1*, which encodes Munc18-1, a synaptic protein essential for synaptic vesicle exocytosis and neurotransmission, and in *SCN2A*, which encodes the voltage-gated sodium channel Nav1.2 (αII subunit), have been identified in patients with epilepsy. Although haploinsufficiency of either *Stxbp1* or *Scn2a* in cortical excitatory neurons induces epileptic phenotypes in mice and a reduction of cortico-striatal, especially cortico-striatal parvalbumin-expressing fast-spiking interneurons (FSIs), excitatory transmission has been suggested to be the basis, the subcortical circuits remain poorly understood. In this study, we investigated which striatal subregions’ FSIs are responsible for the epileptic seizures. Using chemogenetic approach, we selectively suppressed FSI activity in either the nucleus accumbens (NAc) or the dorsal striatum (caudate–putamen, CPu) of mice. Suppression of FSIs in the NAc induced outwardly-recognized convulsive seizures accompanied by epileptiform discharges in the electrocorticographic (ECoG) analysis, whereas inhibition of FSIs in the CPu resulted in epileptiform discharges without overt convulsions. Notably, focal suppression of FSIs in either the anterior or medial region of the NAc shell (NAcSh), but not in other NAc subregions, was sufficient to trigger convulsive seizures. These findings identify FSIs in the anteromedial shell of the NAc as a critical hub for convulsive seizure generation and provide new insights into the striatal circuit mechanisms underlying *STXBP1*- or *SCN2A*-associated epilepsies.

## Introduction

Munc18-1, encoded by the *STXBP1* gene, is an essential presynaptic protein that plays a critical role in synaptic vesicle docking and neurotransmitter release [Dulubova *et al*.,2007]. In the mouse brain, Munc18-1 is widely expressed across neuronal populations, including both excitatory and inhibitory neurons, whereas its expression is largely restricted to neurons and is absent from glial cells [Verhage *et al*., 2000; Toonen *et al*., 2006]. At the subcellular level, Munc18-1 is highly enriched at presynaptic terminals, where it associates with syntaxin-1 and other components of the SNARE complex to regulate synaptic vesicle priming and fusion [Rizo and Südhof, 2012]. The voltage-gated sodium channel Nav1.2, encoded by *SCN2A*, is essential for the initiation and propagation of action potentials in excitatory neurons, thereby shaping cortical network excitability and information processing [Catterall, 1992, Catterall *et al*., 2005]. Nav1.2 is abundantly expressed in the axons of cortical and hippocampal excitatory neurons [Liao *et al*., 2010; Ogiwara *et al*., 2018; Yamagata *et al*., 2023], in dopaminergic neurons of the substantia nigra pars compacta and the ventral tegmental area (VTA) [Yang *et al*., 2019], as well as in specific populations of inhibitory neurons, including medium spiny neurons (MSNs) in the striatum [Miyazaki *et al*., 2014] and GABAergic interneurons in the neocortex [Li *et al*., 2014; Yamagata *et al*., 2017].

Pathogenic mutations in both *STXBP1* and *SCN2A* are known to cause a wide spectrum of neurodevelopmental disorders, including various forms of epilepsy, autism spectrum disorder (ASD), intellectual disability (ID), and schizophrenia [Sugawara *et al*, 2001; Kamiya *et al*, 2004; Saitsu *et al*., 2008; Ogiwara *et al*, 2009; Buxbaum *et al*, 2012; Tavassoli *et al*, 2014; Li *et al*, 2016; Rauch *et al*, 2012; de Ligt *et al*, 2012; Fromer *et al*, 2014; Hoischen *et al*, 2014; Johnson *et al*, 2016; Carroll *et al*, 2016; Stamberger *et al*., 2016; Balakrishna *et al*, 2020; Stamberger *et al*., 2022]. Rodent models with *Scn2a* haploinsufficiency exhibit epileptic phenotypes, cognitive impairments, and behavioral abnormalities reminiscent of *SCN2A*-associated disorders in humans [Ogiwara *et al*., 2018, Middleton *et al*., 2018; Tatsukawa, 2019; Suzuki, 2024]. Our previous studies using genetic models of *STXBP1-* and *SCN2A*-related epilepsies have provided evidence that reduced excitatory transmission from cortical pyramidal neurons onto striatal parvalbumin-expressing (PV^+^) fast-spiking interneurons (FSIs) is sufficient to trigger epilepsy [Miyamoto *et al*., 2019; Ogiwara *et al*., 2018]. Consistently, pharmacological inhibition of cortico-striatal excitatory inputs onto FSIs using Ca²⁺-permeable AMPA receptor antagonists has been shown to evoke generalized seizures in both mice and non-human primates [Gittis *et al*., 2011; Aupy *et al*., 2023]. In contrast, other group has reported that direct optogenetic or chemogenetic silencing of FSIs in the CPu alone fails to reliably induce seizures [Lee *et al*., 2017], suggesting that CPu FSIs may not play a major role in seizure generation. On the other hand, accumulating evidences suggest that the ventral striatum, particularly the nucleus accumbens (NAc), also contributes to seizure modulation. For example, altered activity of MSNs in the NAc has been shown to influence epilepsy: reduced activity of dopamine D1 receptor-expressing (D1R⁺) MSNs or enhanced activity of dopamine D2 receptor-expressing (D2R^+^) MSNs exacerbates absence seizures [Deransart *et al*., 1998], whereas MSN hyperactivity is observed in temporal lobe epilepsy (TLE) models induced by intra-amygdala kainic acid (KA) injection, and suppression of this activity alleviates seizures [Zou *et al*., 2022]. Moreover, PV⁺ interneurons in the NAcSh have been implicated in seizure modulation in TLE models induced by intra-hippocampal KA administration [Jiang *et al*., 2024].

In the present study, we investigated whether reduced activity of FSIs in the NAc or the CPu is sufficient to induce epileptic seizures. We found that mice with suppressed FSI activity in the NAc exhibited convulsive seizures accompanied by epileptiform discharges, whereas suppression of FSIs in the CPu resulted in abnormal ECoG activity without overt convulsions. Furthermore, we found that suppressing FSIs in the anterior and medial NAcSh, but not in other NAc subregions, induced convulsive seizures and epileptiform activity. Together, these findings suggest that FSIs in the NAc, particularly those in the anteromedial NAcSh, play a critical role in convulsive seizure generation, whereas CPu FSIs may be less directly involved.

## Results

### Generation of mice with selective inhibition of FSIs in the NAc or the CPu

Previous studies have reported that PV⁺ interneurons are markedly sparse in the NAcSh compared with the CPu and the NAc core (NAcC) [Castro *et al*., 2019; Marinescu *et al*., 2024]. To confirm this anatomical feature under our experimental conditions, we performed immunohistochemical staining with an anti-PV antibody in 3-month-old C57BL/6J wild-type mice. Consistent with previous reports, PV⁺ interneurons were abundantly distributed in the CPu and the NAcC, whereas only a small number of PV⁺ cells were detected in the NAcSh (**Suppl. Fig. S1**). These observations confirm the sparse distribution of this interneuron subtype in the shell region under our experimental conditions.

To achieve selective inhibition of FSIs in either the NAc or the CPu, we bilaterally injected an adeno-associated virus (AAV) vector encoding a Cre-dependent inhibitory DREADD (AAV5-*EF1a*-DIO-hM4D(Gi)-mCherry) into the target regions of *PV*-Cre driver mice. In this driver line, Cre recombinase is selectively expressed in PV⁺ neurons, thereby enabling region-specific expression of hM4D(Gi) in FSIs. Coronal brain sections were prepared from mice at 4 months of age following AAV injection into either the NAc or the CPu. Histological analyses confirmed accurate targeting of the injection sites. In mice injected in the NAc, mCherry fluorescence was confined to the NAc, with no detectable spread into the CPu (**Fig. 1**). Conversely, in mice injected in the CPu, fluorescence signals were restricted to the CPu, with no apparent leakage into the NAc (**Fig. 1**). Together, these results demonstrate region-specific viral expression, enabling selective inhibition of FSIs in either the NAc or the CPu.

**Figure 1.**
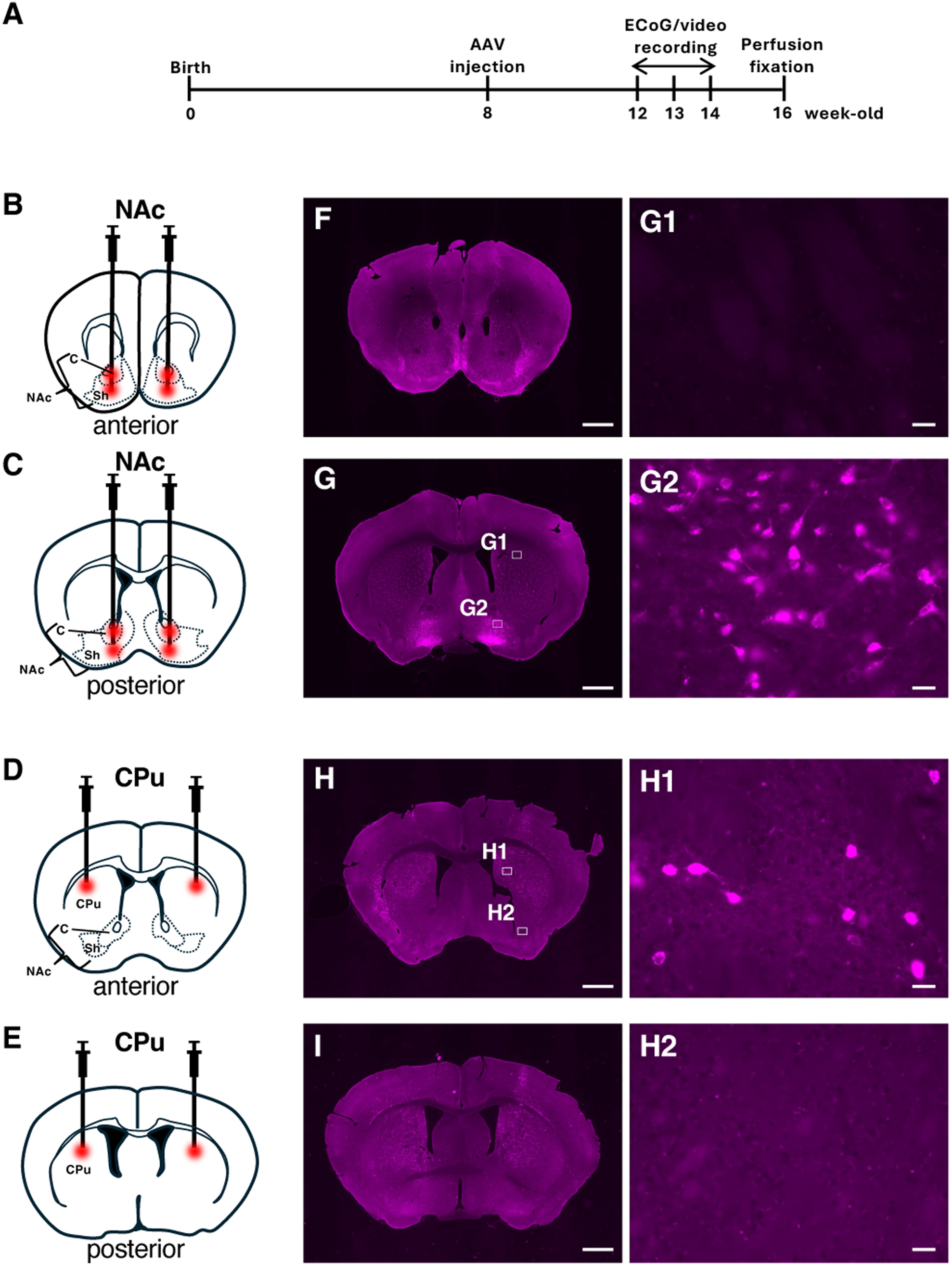
AAVs were selectively injected into either the NAc or the CPu. (**A**) Schematic diagram of the experimental procedure. (**B-E**) Schematic diagrams of illustrating viral injection into the NAc (B, C) or the CPu (D, E) of *PV*-Cre mice. The target regions for viral injection are indicated in red. (**F, G**) In coronal frozen brain sections obtained near the injection sites, mCherry expression was predominantly observed in the NAc. G1 (CPu region) and G2 (NAc region) show higher-magnification images of the areas outlined in G. (**H, I**) Fluorescence signals were restricted to the CPu. H1 (CPu region) and H2 (NAc region) show higher-magnification images of the areas outlined in H. Scale bars: 1 mm (F–I); 20 µm (G1, G2, H1, H2). AAV, adeno-associated virus; C, core; CPu, caudate–putamen; NAc, nucleus accumbens; Sh, shell.

### Suppression of FSIs in the NAc, but not in the CPu, induces convulsive seizures

We next performed simultaneous ECoG recordings and video monitoring in these mice to examine the functional consequences of FSI inhibition in each region. In mice with chemogenetic suppression of FSIs in the NAc, systemic administration of the DREADD agonist clozapine-N-oxide (CNO) induced convulsive seizures accompanied by robust epileptiform discharges (**Suppl. Video S1**). The ECoG traces exhibited high-amplitude, synchronized polyspike activity that temporally correlated with behavioral manifestations, including convulsive seizures. These findings indicate that reduced activity of NAc FSIs is sufficient to trigger convulsive seizure activity. In contrast, inhibition of FSIs in the CPu did not elicit overt convulsive seizures. Although abnormal ECoG activity was observed following chemogenetic suppression in the CPu, these abnormalities did not progress to convulsive seizures (**Suppl. Video S2**). Quantitative analysis further revealed that the percentage of mice exhibiting convulsive seizures was significantly higher in the NAc-FSI–suppressed group than in the CPu-FSI–suppressed group (**Fig. 2A**). In addition, the frequency of abnormal ECoG events was significantly greater in NAc-manipulated mice compared with CPu-manipulated mice (**Fig. 2B**). Because the CPu is substantially larger in volume than the NAc, we further examined whether increasing the extent of viral transduction in the CPu would alter the outcome. To this end, we doubled the volume of AAV injected into the CPu and performed the same chemogenetic suppression and ECoG analyses. Even under these conditions, mice exhibited only abnormal ECoG activity without progression to overt convulsive seizures, similar to the original CPu-manipulated group (**Fig. 2A, B**). Together, these results suggest that FSIs in the NAc play a more prominent role in the generation of convulsive seizures than FSIs in the CPu.

**Figure 2.**
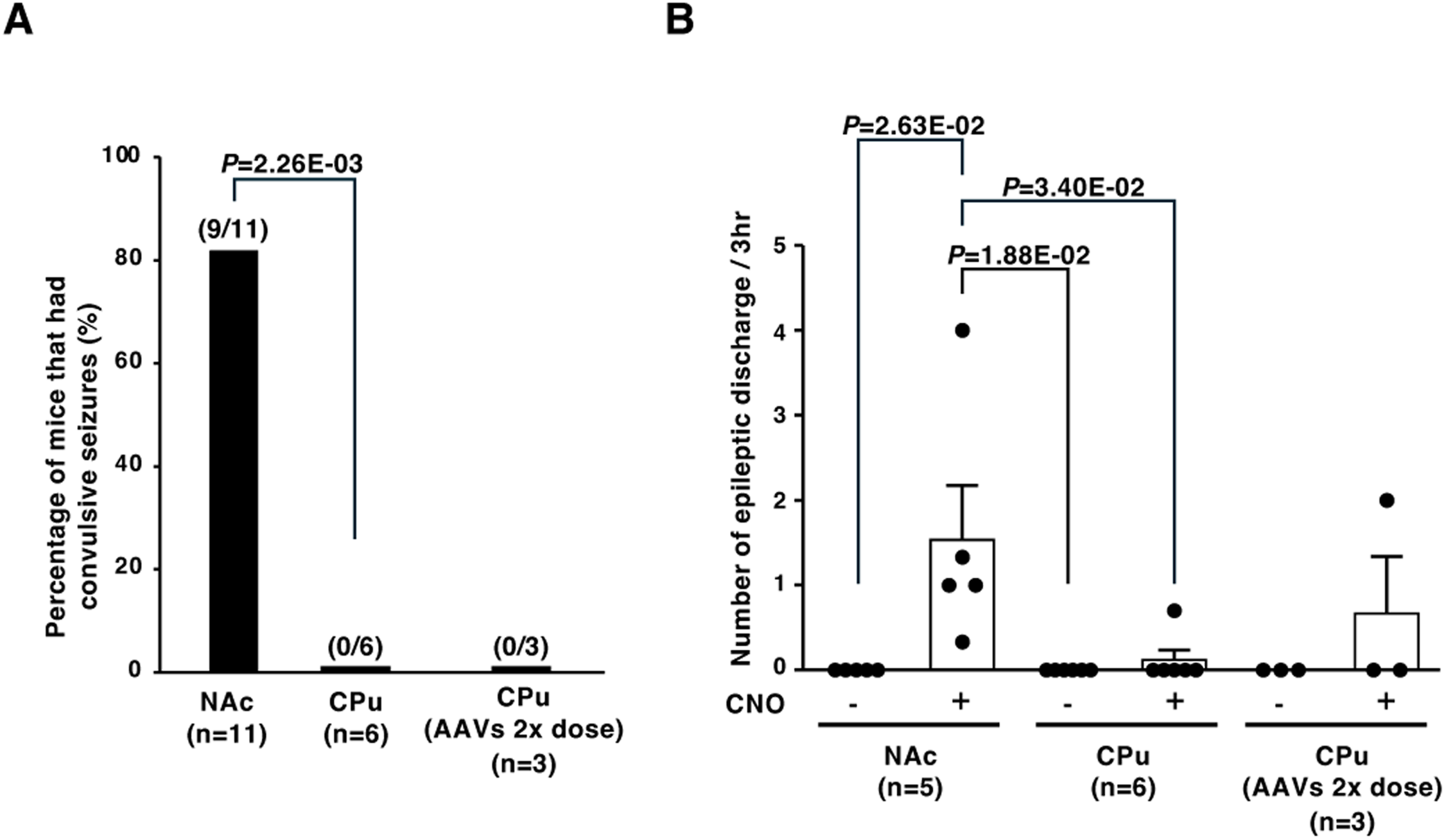
FSI suppression in the NAc, but not in the CPu, induces convulsive seizures. (**A**) Incidence of convulsive seizures following FSI suppression in either the NAc or the CPu of *PV*-Cre mice (3-h observation period after CNO injection). (**B**) Number of epileptiform discharges in mice with FSI suppression in either the NAc or the CPu (3-h recordings before [−] and after [+] CNO injection). Black dots represent individual mice. The number of mice is indicated in parentheses. AAV, adeno-associated virus; CNO, clozapine-N-oxide; CPu, caudate–putamen; NAc, nucleus accumbens.

### Suppression of FSIs in the anterior or medial NAcSh, but not in other NAc subregions, induces convulsive seizures

To further delineate which NAc subregion is responsible for seizure generation, we generated additional cohorts of *PV*-Cre mice in which the Cre-dependent inhibitory DREADD was bilaterally injected into specific NAc subregions (**Fig. 3A-F**). Targeted injections were made into the anterior (rostral), posterior (caudal), medial, or lateral portions of the NAcSh, as well as into the anterior or posterior NAcC. Following recovery, simultaneous ECoG recordings and video monitoring were performed as described above. Remarkably, suppression of FSIs in the anterior shell or medial shell resulted in convulsive seizures accompanied by epileptiform ECoG abnormalities in one of two mice per group (**Fig. 3G, H**). In these animals, high-amplitude synchronized discharges were temporally associated with behavioral convulsive episodes (**Suppl. Videos S3 and S4)**. In contrast, inhibition of FSIs in the posterior shell, lateral shell, or either anterior or posterior core did not induce convulsive seizures in any of the examined mice (**Fig. 3 G, H**). These findings narrow the seizure-relevant region within the NAc and suggest that FSIs located in the anteromedial shell play a particularly prominent role in the generation of convulsive seizures.

**Figure 3.**
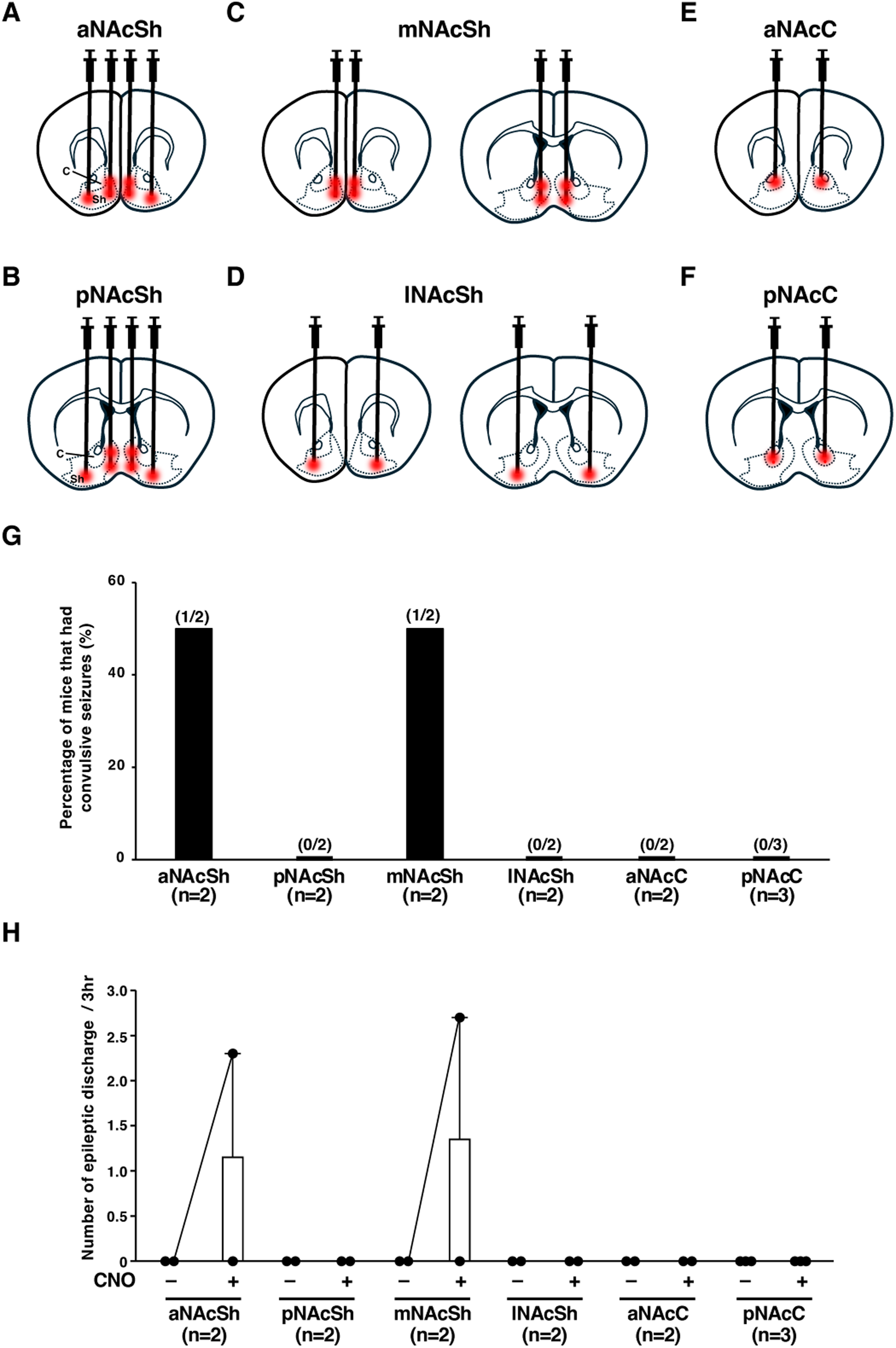
FSI suppression in the anteromedial NAcSh, but not in other NAc subregions, induces convulsive seizures. (**A-F**) Schematic diagrams illustrating viral injections into specific NAc subregions in *PV*-Cre mice. The target regions for viral injection are indicated in red. (**G**) Incidence of convulsive seizures following FSI suppression in specific NAc subregions of *PV*-Cre mice (3-h observation period after CNO injection). (**H**) Number of epileptiform discharges in mice with FSI suppression in specific NAc subregions (3-h recordings before [−] and after [+] CNO injection). Black dots represent individual mice. The number of mice is indicated in parentheses. aNAcC, anterior NAc core; aNAcSh, anterior NAc shell; C, core; CNO, clozapine-N-oxide; lNAcSh, lateral NAc shell; mNAcSh, medial NAc shell; NAc, nucleus accumbens; pNAcC, posterior NAc core; pNAcSh, posterior NAc shell; Sh, shell.

## Discussion

In this study, we investigated the epileptic phenotypes of mice with selective inhibition of FSIs in either the NAc or the CPu. We found that inhibition of FSIs in the NAc, but not in the CPu, induced convulsive seizures and epileptiform activity.

In our previous study, we reported that manipulation of FSIs in the CPu could either induce or suppress spike-and-wave discharges (SWDs) characteristic of absence seizures, and that inhibition of excitatory inputs to CPu FSIs causes a gradual progression from mild to convulsive seizures in a dose-dependent manner [Miyamoto *et al*., 2019]. In the present study, we found that reduced activity of FSIs in the CPu, which are primarily involved in motor control, was associated only with minor epileptiform discharges without convulsive seizures. Unexpectedly, we discovered that reduced activity of FSIs in the NAc, a region implicated in reward and emotion, was sufficient to generate convulsive seizures. These results suggest that NAc FSIs constitute a critical component of the neural circuitry underlying convulsive seizure generation, whereas CPu FSIs may exert a more modulatory or indirect influence.

The NAc is anatomically and functionally divided into two subregions: the core, which is involved in the acquisition of reward–cue associations and the initiation of motor actions, and the shell, which is associated with reward prediction and affective processing [Klawonn *et al*., 2018]. Here, we found that selective inhibition of FSIs in the anterior part of the NAcSh, possibly the anteromedial shell, induced convulsive seizures. Although previous studies have suggested that the NAc may play a role in seizure modulation, most of these reports were based on pharmacologically induced TLE mouse models [Zou *et al*., 2022; Jiang *et al*., 2024]. To our knowledge, this is the first report showing that inhibition of NAcSh FSIs alone, without the use of convulsant agents, can induce spontaneous convulsive seizures. This finding expands the current understanding of NAc function by identifying a previously unrecognized role in ictogenesis. Interestingly, despite the fact that the NAcSh contains fewer PV^+^ FSIs compared to the CPu or NAcC, suppression of this relatively small population was sufficient to elicit convulsive seizures. However, it remains unclear how decreased activity of FSIs in the NAc, particularly in the shell region, which is more closely associated with motivational and emotional processes than with motor control, can lead to convulsive seizures accompanied by overt motor manifestations. It also remains possible, though unlikely, that inadvertent viral leakage to surrounding regions, such as the olfactory tubercle or ventrolateral cortex, reduced PV^+^ neuronal activity and played a role in triggering seizures.

Taken together with previously reported neural circuitries [Kupchik *et al*., 2015; Qi *et al*. 2016; Chen *et al*., 2019; Miyamoto *et al*, 2019; Soares-Cunha *et al*., 2023; Suzuki *et al*., 2024; Xu *et al*, 2024; Marinescu *et al*., 2024], we propose that a circuit comprising the NAcSh, VP, thalamus, and neocortex may contribute to convulsive seizure generation (**Fig. 4**). In this model, impaired activity of FSIs in the NAcSh leads to disinhibition of D2R^+^ MSNs. Hyperactivation of these D2R^+^ MSNs would result in excessive inhibition of VP neurons, which are primarily PV^+^ cells. Such suppression of VP activity may reduce inhibitory synaptic input to the thalamus, thereby promoting abnormal thalamocortical excitation and ultimately facilitating convulsive seizures. However, several studies investigating NAc microcircuits in the context of aversion and preference—rather than epilepsy—have reported findings that appear inconsistent with this framework. For example, Qi and colleagues [Qi *et al*. 2016] demonstrated that glutamatergic inputs from the VTA to the NAc drive aversion by acting on GABAergic interneurons, and that direct photoactivation of NAc PV^+^ interneurons was sufficient to elicit aversive behavior. In contrast, Chen and colleagues [Chen *et al*., 2019] reported that activation of accumbal PV^+^ interneurons induced place preference, whereas their inhibition resulted in conditioned place aversion. Moreover, in the context of epilepsy, the present hypothesis appears to contradict our previous observation that excessive inhibition of thalamic neurons can trigger SWDs and epileptogenesis. This apparent discrepancy raises the possibility that striatal inputs from the CPu and the NAc reach the thalamus through distinct intermediary nuclei, such as the globus pallidus interna/substantia nigra pars reticulata or VP and terminate in different thalamic subregions. Importantly, regional differences in the expression of Cav3.1 T-type calcium channels, which mediate rebound bursting in thalamic neurons [Kim *et al*., 2001], may critically influence whether altered thalamic inhibition suppresses or facilitates seizure activity. Further investigation will therefore be required to define how modulation of thalamic inhibition leads to distinct seizure phenotypes. Overall, this pathway may represent a previously unrecognized mechanism linking the limbic system, particularly the NAc, which is strongly associated with reward processing and emotional regulation, to motor networks. Such a mechanism provides new insight into the subcortical control of convulsive seizure generation.

**Figure 4.**
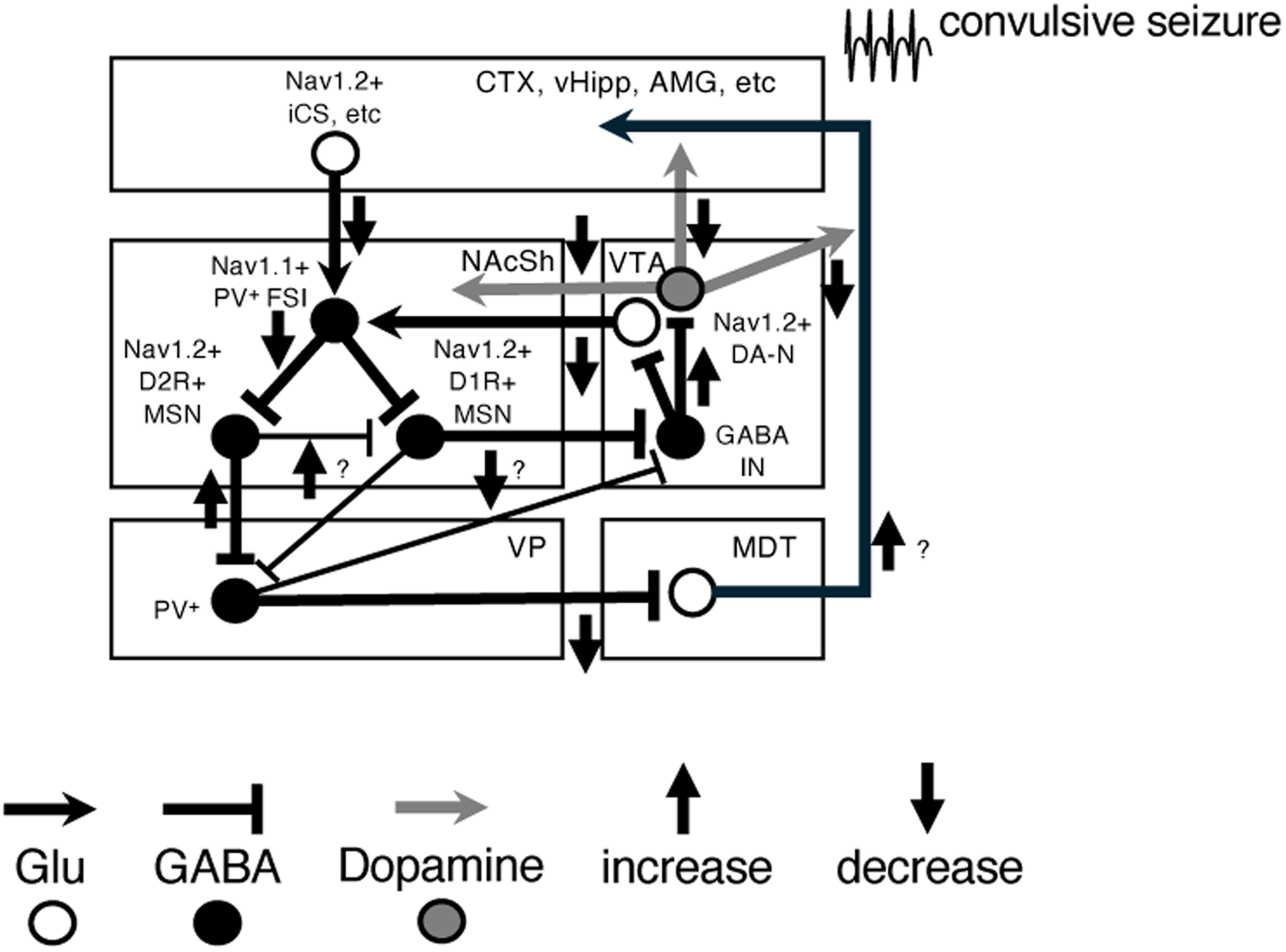
Circuit model for convulsive seizure generation in mice with impaired FSI activity in the NAcSh. This schematic model of the neural circuit was modified and simplified from previously published diagrams (Kupchik *et al*., 2015; Qi *et al*., 2016; Chen *et al*., 2019; Miyamoto *et al*., 2019; Soares-Cunha *et al*., 2023; Suzuki *et al*., 2024; Xu *et al*., 2024; Marinescu *et al*., 2024). Upward and downward arrows indicate increased and decreased neural transmission, respectively. AMG, amygdala; CTX, neocortex; DA-N, dopaminergic neuron (gray circle) and transmission (gray arrows); FSI, fast-spiking interneuron; GABA, GABAergic neurons (black circles) and transmission (black lines); Glu, glutamatergic neurons (white circles) and transmission (black arrows); iCS, intratelencephalic cortico-striatal neurons; IN, interneurons; MDT, mediodorsal thalamus; MSN, medium spiny neuron; NAcSh, nucleus accumbens shell; Nav1.1, voltage-gated sodium channel α1 subunit; Nav1.2, voltage-gated sodium channel α2 subunit; vHippo, ventral hippocampus; VP, ventral pallidum; VTA, ventral tegmental area.

In summary, our results showed that inactivation of FSIs in the NAc contributes to convulsive seizure generation, whereas inactivation of FSIs in the CPu results only in abnormal ECoG activity without overt convulsions. These findings advance our understanding of the pathological neural circuitry underlying convulsive seizures in epilepsy.

## Methods

### Animals

Parvalbumin (PV)-Cre transgenic mice [Tanahira *et al*., 2009] (C57BL/6 J background) were maintained on 12 h light/dark cycle with ad libitum access to food and water. Genotyping of mouse was performed by PCR analysis using following primer pair: Cre-1F: 5’–cgaacgcactgatttcgacc–3’ and Cre-1R: 5’– aaccagcgttttcgttctgc–3’ (product size = Cre allele 202 bp).

### AAVs

Following AAV vectors were used in this study, with microinjection as indicated. Plasmid construct for double floxed Gi-coupled hM4D DREADD fused with mCherry under the control of EF1a promoter, pAAV-EF1a-DIO-hM4D(Gi)-mCherry was a gift from Dr. Bryan Roth (Addgene plasmid 50461). Packaging of the plasmid vector into AAVs, purification, and quantification of the viruses were performed at the Division of Genetic Therapeutics of the Jichi Medical University and at the section for viral vector development of the National Institute of Physiological Sciences.

### Stereotaxic surgery

PV-Cre transgenic mice (>8 weeks of age, both sexes) were anesthetized with isoflurane (1.0–2.5%) and placed in a stereotaxic apparatus (Stoelting). Bilateral injections of AAV5-EF1α-DIO-hM4D(Gi)-mCherry (7.5 × 10⁹ viral genomes/μL) or phosphate-buffered saline (PBS) were performed using a microinjector (Nanoliter 2020 Injector; World Precision Instruments) fitted with a pulled glass capillary at a flow rate of 100 nL/min. Stereotaxic coordinates were determined according to the mouse brain atlas [Paxinos *et al*., 2001]. Detailed microinjection coordinates and injection volumes are provided in **Supplementary Table S1**.

### Electrocorticogram (ECoG) recording

Three weeks after AAV vector injection, mice (>11 weeks of age) were used for experiments. ECoG was performed to directly monitor cortical activity using stainless steel screw electrodes implanted in the skull. Stainless steel screws (1.1 mm in diameter) serving as ECoG electrodes were placed bilaterally over the somatosensory cortex (±1.5 mm lateral to the midline and 1.0 mm posterior to bregma) under 1.0–2.5% isoflurane anesthesia. Two stainless steel screw serving as the reference and ground electrode were placed over the cerebellum. For electromyography (EMG), a bipolar stainless steel wire electrode (100 μm diameter) was inserted into the trapezius muscle in the cervical region. Recordings were initiated at least one week after surgery and at least four weeks after AAV injection. ECoG and infrared video recordings were performed for 3 h before clozapine-N-oxide (CNO) administration. CNO (2 mg/kg, dissolved in 0.5% DMSO in saline) was administered intraperitoneally, and recordings were continued for 3 h starting 30 min after injection. The frequency of ECoG abnormalities was quantified by averaging three recording sessions per mouse, each separated by approximately one week.

### Histological analyses

Mice were deeply anesthetized and transcardially perfused with 0.9% NaCl, followed by 4% paraformaldehyde (PFA) in 0.1 M phosphate buffer (PB). Brains were removed and post-fixed in 4% PFA for 36 h at 4°C, followed by cryoprotection in 30% sucrose in 0.1 M PB for 36 h at 4°C. The brains were then embedded in Tissue-Tek O.C.T. compound (Sakura Finetek). Coronal brain sections (30 μm thick) were prepared using a cryostat (Leica CM1900; Leica Biosystems). Immunohistochemical analyses, 6-μm-thick paraffin sections prepared from 4% periodate–lysine–paraformaldehyde (PLP)-fixed mouse brains were processed as described previously [Yamagata *et al*., 2023]. The sections were incubated with a mouse anti-PV monoclonal antibody (Swant, 1:500) for 12 h at 4 °C. After washing three times with PBS, sections were incubated with Alexa Fluor™ Plus 555‒conjugated anti-mouse IgG secondary antibody (Thermo Fisher Scientific, 1:1,000) for 1 h at room temperature. Images were acquired using a BZ-X710 fluorescence microscope (Keyence).

### Statistical analyses

Statistical significance was assessed using one-way analysis of variance (ANOVA) followed by Tukey’s post hoc test (KyPlot; KyensLab Inc.) and 2 × 3 chi-square tests using Pearson’s method. For the chi-square tests, a Bonferroni correction was applied to adjust for multiple comparisons, resulting in a corrected significance level of 0.017. Data are presented as the mean ± standard error of the mean (SEM), and statistical significance was defined as p < 0.05.

## Acknowledgments

The authors thank Drs. Tamamaki and Tanahira (Kumamoto University) for generously providing the PV-Cre-TG driver line. We also thank the members of the Department of Neurodevelopmental Disorder Genetics and the staff of the animal facility at Nagoya City University (NCU) for their support. We are grateful to the Research Equipment Sharing Center at NCU for technical assistance.

## Funding

This work was supported by grants from NCU; JSPS KAKENHI (Grant Numbers 23K27490 and 25K10820); and the Grant-in-Aid for Outstanding Research Group Support Program at NCU (Grant Number 2401101). This work was also supported by the use of research equipment shared under the MEXT Project for Promoting Public Utilization of Advanced Research Infrastructure (Program for Supporting Construction of Core Facilities) (Grant Number JPMXS0441500024).

## Data Availability

The datasets generated during and/or analyzed during the current study are available from the corresponding author on reasonable request.

## Declarations

### Ethics Approval

All animal breeding and experimental procedures were approved by the Institutional Animal Care and Use Committee of NCU. All procedures were conducted in accordance with the ARRIVE guidelines and the institutional guidelines and regulations of NCU.

### Consent to Participate

Not applicable.

### Consent for Publication

Not applicable.

### Competing Interests

The authors declare no competing interests.

## Notes

### Competing Interest Statement

The authors have declared no competing interest.

