## Supplementary Table S1 for "Reduced activity of nucleus accumbens parvalbumin-expressing fast-spiking inhibitory neurons causes convulsive seizures"

| Injection region | AP | ML | DV | Injection volume / site |
|---|---|---|---|---|
| **NAc** | +1.60 | ±1.00 | -5.00 | 250 nl |
| | +1.60 | ±1.00 | -4.50 | |
| | +0.80 | ±1.00 | -5.20 | |
| | +0.80 | ±1.00 | -4.50 | |
| **CPu** | +0.70 | ±2.00 | -3.00 | 500 nl |
| | +0.00 | ±2.00 | -2.80 | |
| **CPu (AAVs 2x dose)** | +0.70 | ±2.00 | -3.50 | 500 nl |
| | +0.70 | ±2.00 | -3.00 | |
| | +0.00 | ±2.00 | -3.50 | |
| | +0.00 | ±2.00 | -2.80 | |
| **aNAcSh** | +1.60 | ±0.45 | -4.80 | 250 nl |
| | +1.60 | ±0.45 | -4.25 | |
| | +1.60 | ±1.20 | -4.90 | |
| **pNAcSh** | +0.80 | ±0.54 | -4.80 | 250 nl |
| | +0.80 | ±0.54 | -4.30 | |
| | +0.80 | ±1.15 | -5.20 | |
| **mNAcSh** | +1.60 | ±0.45 | -4.80 | 100 nl |
| | +1.60 | ±0.45 | -4.25 | |
| | +0.80 | ±0.54 | -4.80 | |
| | +0.80 | ±0.54 | -4.30 | |
| **lNAcSh** | +1.60 | ±1.20 | -4.90 | 100 nl |
| | +0.80 | ±1.15 | -5.20 | |
| **aNAcC** | +1.60 | ±0.90 | -4.40 | 100 nl |
| **pNAcC** | +0.80 | ±1.00 | -4.70 | 100 nl |

**Table S1. AAV microinjection coordinates and injection volume.**
NAc, nucleus accumbens; CPu, Caudate Putamen; aNAcSh, anterior NAc shell; pNAcSh, posterior NAc shell; mNAcSh, medial NAc shell; lNAcSh, lateral NAc shell; aNAcC, anterior NAc core; pNAcC, posterior NAc core; AP, anteroposterior; ML, mediolateral; DV, dorsoventral.
