## Supplementary figures and images for "Reduced activity of nucleus accumbens parvalbumin-expressing fast-spiking inhibitory neurons causes convulsive seizures"

### Supplementary Figure S1

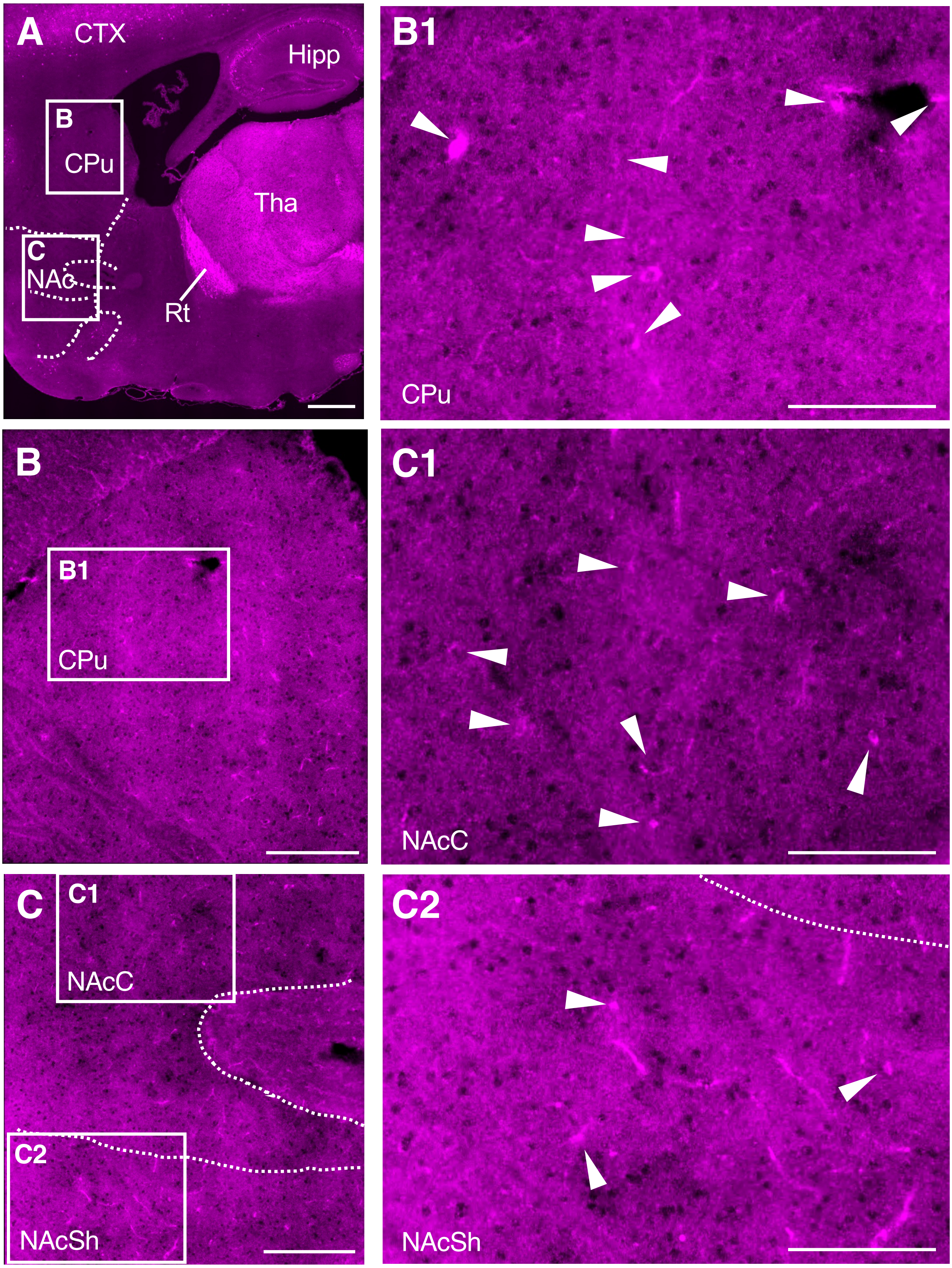
